# Landscape and regulation of m^6^A and m^6^Am methylome across human and mouse tissues

**DOI:** 10.1101/632000

**Authors:** Jun’e Liu, Kai Li, Jiabin Cai, Mingchang Zhang, Xiaoting Zhang, Xushen Xiong, Haowei Meng, Xizhan Xu, Zhibin Huang, Jia Fan, Chengqi Yi

## Abstract

*N*^6^-methyladenosine (m^6^A), the most abundant internal mRNA modification, and *N*^6^,2’-O-dimethyladenosine (m^6^Am), found at the first-transcribed nucleotide, are two examples of dynamic and reversible epitranscriptomic marks. However, the profiles and distribution patterns of m^6^A and m^6^Am across different human and mouse tissues are poorly characterized. Here we report the m^6^A and m^6^Am methylome through an extensive profiling of 42 human tissues and 16 mouse tissue samples. Globally, the m^6^A and m^6^Am peaks in non-brain tissues demonstrates mild tissue-specificity but are correlated in general, whereas the m^6^A and m^6^Am methylomes of brain tissues are clearly resolved from the non-brain tissues. Nevertheless, we identified a small subset of tissue-specific m^6^A peaks that can readily classify the tissue types. The number of m^6^A and m^6^Am peaks are partially correlated with the expression levels of their writers and erasers. In addition, the m^6^A- and m^6^Am-containing regions are enriched for single nucleotide polymorphisms. Furthermore, cross-species analysis of m^6^A and m^6^Am methylomes revealed that species, rather than tissue types, is the primary determinant of methylation. Collectively, our study provides an in-depth resource for dissecting the landscape and regulation of the m^6^A and m^6^Am epitranscriptomic marks across mammalian tissues.

## INTRODUCTION

More than 100 RNA modifications have been characterized so far (Machnicka et al., 2013). *N*^6^-methyladenosine (m^6^A) is the most prevalent post-transcriptional modification of messenger RNA (mRNA) and long noncoding RNA (lncRNA) in mammalian cells (Li et al., 2016; Liu and Pan, 2016; Zhao et al., 2017). m^6^A modification is catalyzed by a multi-component methyltransferase complex that consists at least of METTL3, METTL14, WTAP, KIAA1429 and RBM15 (Bokar et al., 1994; Bokar et al., 1997; Liu et al., 2014; Ping et al., 2014; Schwartz et al., 2014; Wang et al., 2014b). As the first reversible mRNA modification, m^6^A can be “erased” via FTO and ALKBH5 (Jia et al., 2011; Zheng et al., 2013). In addition, m^6^A can be recognized by different types of reader proteins: the YTH-family proteins that specifically recognize the m^6^A modification in RNA, proteins with a common RNA binding domain and its flanking regions together to bind m^6^A, and proteins that use a so-called m^6^A-switch mechanism for recognition (Hsu et al., 2017; Roundtree et al., 2017b; Shi et al., 2017; Wang et al., 2014a; Wang et al., 2015; Zhou and Pan, 2018). These reader proteins further lead to different biological outcomes for the m^6^A-marked RNA transcripts.

Dynamic and reversible m^6^A has been shown to play critical roles in RNA metabolism, physiological and pathological processes. For instance, m^6^A is involved in RNA splicing, export, stability, translation and localization (Roundtree et al., 2017a; Yang et al., 2018). Moreover, m^6^A-dependent mRNA regulation has now been demonstrated to regulate the process of circadian rhythm, adipogenesis, spermatogenesis, embryonic stem cell self-renewal and differentiation (Fu et al., 2014). The aberrant regulation of m^6^A is related to a variety of cancers including acute myeloid leukemia, breast cancer, glioblastoma, lung cancer and liver cancer (Hong, 2018; Luo et al., 2018), which highlight the important regulatory roles of m^6^A.

Different from the internal m^6^A, there exists a terminal modification, termed as *N*^6^,2’-*O*-dimethyladenosine (m^6^Am). m^6^Am was originally discovered at the 5’ end of mRNA in animal cells and viruses in 1975 (Wei et al., 1975). The 2’-hydroxyl position of the ribose sugar of the first and often second nucleotide following the *N*^7^- methylguanosine (m^7^G) mRNA cap can be methylated by 2′-*O*-methyltransferases (2′-*O*-MTases), termed CMTr1 and CMTr2 (Belanger et al., 2010; Werner et al., 2011). If the first nucleotide adjacent to the m^7^G cap is 2′-*O*-methyladenosine (Am), it could be further methylated at the *N*^6^-position to form m^6^Am (Figure 1A). It was only recently discovered that m^6^Am is also reversible by FTO (Mauer et al., 2017; Wei et al., 2018). Moreover, we and others discovered that phosphorylated CTD-interacting factor 1 (PCIF1) is the cap-specific, terminal *N*^6^-methylation enzyme for m^6^Am (Akichika et al., 2018; Sun et al., 2018). While FTO works both on terminal m^6^Am and internal m^6^A, the m^6^Am writer PCIF1 is specific for the terminal m^6^Am modification and hence will further promote the study of the biological functions of m^6^Am in near future.

**Figure 1.**
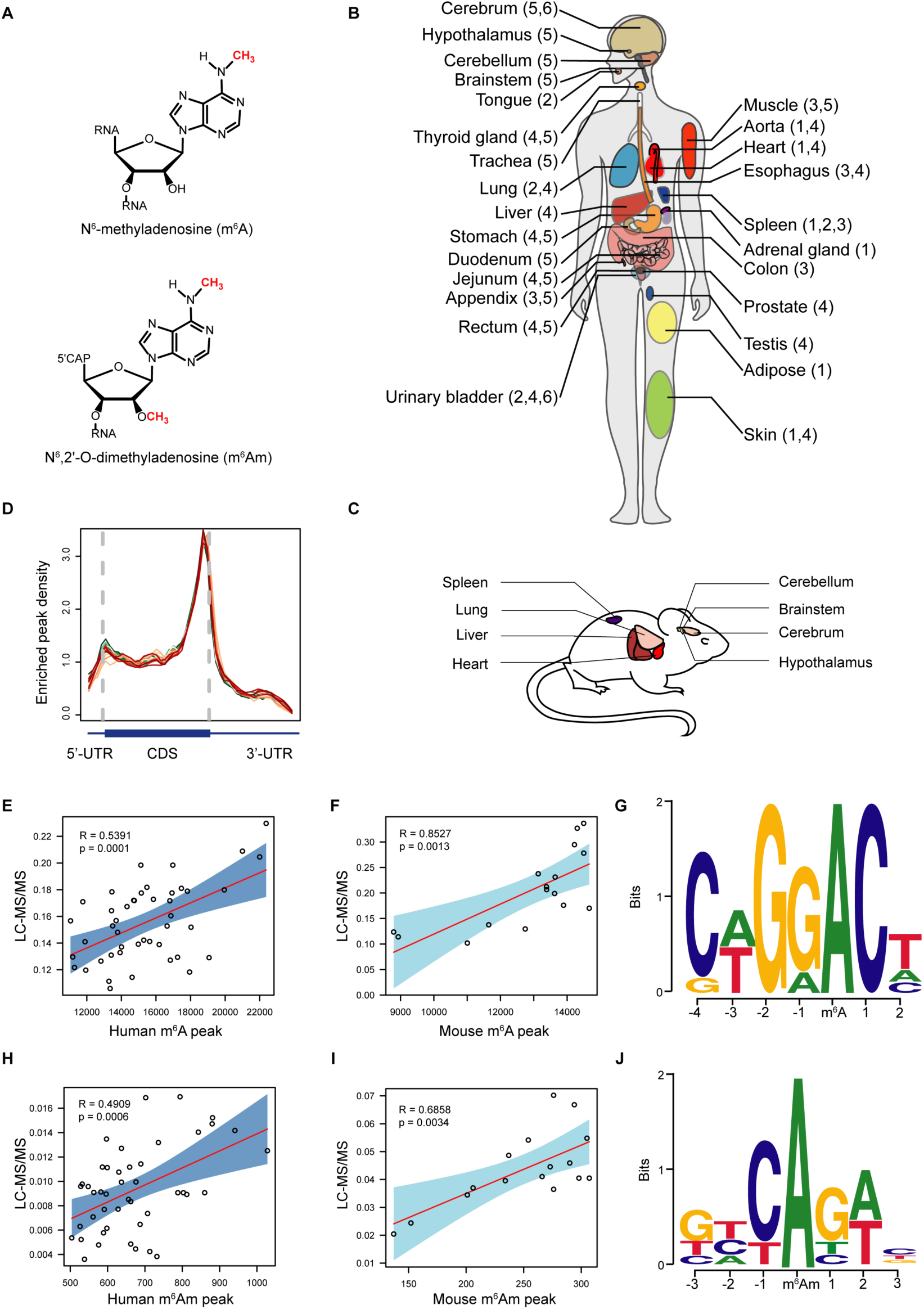
Overviews of m^6^A and m^6^Am in the collected samples of human and mouse tissues. (**A**) Chemical structures of m^6^A and m^6^Am. (**B**) Human tissues analyzed in this study. Tissues collected from donors are denoted and values in parentheses represent the donor IDs. (**C**) Mouse tissues analyzed in this study. (**D**) Distribution of the enriched m^6^A peaks in all human tissues analyzed along the mRNA segments. Each segment was normalized according to its average length in Refseq annotation. Each line denotes the m^6^A distribution in one sample. (**E-F**) Scatter plots showing the correlations between the m^6^A abundance and the m^6^A peak numbers across human (**E**) and mouse tissues (**F**). (**G**) Motif analysis revealed a GRACH consensus for m^6^A peaks in human cerebellum (E-value= 5.5e^-46^). Similar motifs are also observed for m^6^A peaks in other tissue samples. (**H-I**) Scatter plots showing the correlations between the m^6^Am abundance and the m^6^Am peak numbers across human (**H**) and mouse tissues (**I**). (**J**) Motif analysis revealed a BCA consensus for m^6^Am peaks in cerebellum (E-value= 1.6e-^45^). Similar motifs are also observed for m^6^Am peaks in other tissue samples as well.

Several epigenetic and epitranscriptomic marks have been systematically profiled at the tissue and cell level. For instance, the methylome of 5-methylcytosine in DNA (5mdC) across diverse cell lines and tissues revealed that the majority (nearly 80%) of CpG sites are similarly methylated; yet, using the ∼20% differentially methylated regions (DMRs), tissue types and putative regulatory CpGs can be effectively classified and captured (Schultz et al., 2015; Ziller et al., 2013). More recently, the dynamic spatial and temporal patterns of inosine have been demonstrated across many tissues of human and other mammals (Tan et al., 2017). It is reported that the overall editing levels are relatively similar across tissues whereas editing levels in non-repetitive coding regions vary. However, m^6^A and m^6^Am methylome at the human tissue level remains unexplored so far. Previous m^6^A methylome was primarily obtained from limited numbers of mammalian cell lines and a few mouse tissues, while newly discovered reversible m^6^Am methylome was even less well characterized.

In this study, we performed comprehensive analyses of m^6^A and m^6^Am methylome in 42 human and 16 mouse tissue samples. We first recapitulated the similar distribution pattern and consensus motif for m^6^A and m^6^Am in human tissues, as those found in cell lines. We then identified “conserved” and “non-conserved” m^6^A peaks and revealed that m^6^A & m^6^Am peaks are correlated across various non-brain tissues at the overall level, whereas methylome of brain tissues show higher tissue-specificity. Interestingly, the tissue-specific m^6^A peaks could readily distinguish different types of human and mouse tissue. By analyzing the association between the peak numbers of m^6^A & m^6^Am and the expression of the known methyltransferases & demethylases, we uncover that m^6^A and m^6^Am are associated with its writers and erasers. Moreover, the m^6^A- and m^6^Am-containing regions are enriched for single nucleotide polymorphisms (SNPs) in all tissues. To gain further insight into m^6^A and m^6^Am variance, we performed cross-species analysis and revealed that species are more determinant than tissue types. Collectively, our study provides in-depth resource toward elucidating the dynamic patterns and regulation of the epitranscriptomic marks m^6^A and m^6^Am across various human and mouse tissues.

## RESULTS

### m^6^A and m^6^Am abundance across human and mouse tissues

To determine the abundance of m^6^A and m^6^Am in different human and mouse tissues, post-mortem samples which include 42 tissues from 6 individuals (5 males and 1 female) and 16 tissues from 2 male mice were subjected to quantitative MS analysis (Figures 1A-1C and S1A –S1E). The quantification of m^6^A follows the standard procedure (Jia et al., 2011), a decapping step was adapted for m^6^Am owning to its association with the mRNA cap (Mauer et al., 2017; Wei et al., 2018). Application of the quantitative MS studies to various human tissues revealed that the m^6^A/A ratio of total RNA is approximately 0.11%∼0.23% (Figure S1D), while the m^6^Am/A ratio ranges from 0.0036% to 0.0169% (Figure S1E). Hence, the variation of m^6^Am content across different tissues seems to be greater than that of m^6^A. Comparatively speaking, the level of m^6^Am is ∼2.2%-11.4% of that of m^6^A in the same tissue. Overall, the results show that both the m^6^A and m^6^Am are widespread in various human and mouse tissues.

### Transcriptome-wide m^6^A and m^6^Am profiling in human and mouse tissues

To investigate the distribution and dynamics of the m^6^A and m^6^Am among different tissues, transcriptome-wide m^6^A and m^6^Am mapping was performed. Due to the limited amount and partial degraded nature of human tissue RNA samples, we adopted a recently published refined RIP-seq protocol for low-input materials with several modifications (Zeng et al., 2018). It is worth mentioning that instead of random priming, a template-switching reaction at 5’ end of RNA template was used during library construction so that the 5’ end sequence information of RNA was preserved. Combining with the existing knowledge of the cap-adjacent position of m^6^Am modification, m^6^Am sites of mRNA can hence be more accurately determined. Moreover, we removed the cDNA originated from rRNA after the reverse transcription instead of performing a poly(A)+ selection, thereby allowing the preservation and profiling of the non-coding RNAs, including long non-coding RNA (lncRNA) and small nuclear RNA (snRNA) in our study. For instance, U2 and U6 snRNAs are enriched across all tissue samples in our study (Figures S1F and S1G), which is consistent with the previous findings that U6 contains a m^6^A43 and U2 contain a m^6^Am at position 30 (Bohnsack and Sloan, 2018). Furthermore, the high correlation between HEK293T samples from different batches of experiments suggests good reproducibility of our epitranscriptomic sequencing (Figure S1H).

We identified 11,060-24,259 m^6^A peaks for the different human and mouse tissues involved in our study, and calculated the distribution pattern of m^6^A in their mRNAs (Table S1 and S2). Consistent with previous observations, we found that m^6^A peaks were markedly enriched near the stop codon and the distribution pattern of m^6^A is similar among all human tissues (Figure 1D). Moreover, the peak numbers are positively correlated with the m^6^A abundance detected by MS in human and mouse (Figures 1E and 1F). To assess the sequence features of m^6^A, we performed motif search of m^6^A-enriched regions. The previously reported consensus “GRACH” motif (where R represents G or A; H represents A, C, or U) were identified in all tissues (Figure 1G).

We also analyzed m^6^Am peaks by distinguishing it from m^6^A peaks, based on detecting methylated transcriptional start sites as previously suggested (Schwartz et al., 2014). We observed a positive correlation of m^6^Am signal with the m^6^Am abundance detected by MS in human and mouse tissues (Figures 1H and 1I). Moreover, we found that the number of m^6^Am peaks in mRNA varies greatly in different samples, ranging from 526 to 1,028 peaks (Table S1). We next analyzed the m^6^Am consensus and found that they are enriched in the canonical BCA motif (A=m^6^Am, B=C, G or U) (Figure 1J). As expected, the nucleotides adjacent to the BCA motif are pyrimidine-rich sequences known to be around TSS (Carninci et al., 2006; Frith et al., 2008; Ni et al., 2010; Plessy et al., 2010).

### Conservation and tissue-specificity of m^6^A across human and mouse tissues

To investigate potential tissue-specificity of the m^6^A methylome, we first classified all the genes into five categories based on their expression across 42 human tissues as previously published (Uhlen et al., 2015) (see Method Details). We next assessed the proportion of m^6^A modified genes and peak intensity among human tissues. Notably, both the percent of modified genes and peak intensity in ubiquitously expressed genes are significantly higher than that of tissue enriched group for human (*P* < 2.2*e^-16^) (Figures S2A and 2B). We also ruled out difference in expression levels as the explanation, since no significant difference was observed for the two categories (Figure S2C). Take the brain related tissues (cerebellum, cerebrum, hypothalamus and brainstem) as examples, the intensity of brain enriched genes is significantly lower than that of brain non-enriched genes (Figure S2D). Collectively, the results imply that ubiquitously expressed genes are more likely to be m^6^A regulated, while tissue enriched genes are more prone to be regulated at transcript levels.

To further explore the tissue-specificity of m^6^A without the interference of gene expression, we compared the m^6^A methylome using the ubiquitously expressed genes across diverse human and mouse tissues. Overall, four clusters were readily found in human: group 1 are methylome signals from brain tissues (cerebellum, cerebrum, hypothalamus and brainstem), group 2 are methylome of rectum and jejunum, group 3 are methylome of other non-brain tissues used in our study, and group 4 are methylome signals of HEK293T cells. This observation suggests that the m^6^A methylome of brain related tissues are highly specific, while the m^6^A methylome of non-brain tissues show certain tissue-specificity (for instance, rectum and jejunum) but are grouped together in general (Figure 2A). This finding is further corroborated with the m^6^A methylome of mouse tissues (Figure 2B). Thus, our observation implies that m^6^A is involved in the regulation of brain-specific functions that are different from the rest of tissues both in human and mouse. Note that there is less variation among mouse tissues; hence a relatively higher tissue-specificity among non-brain tissues is also observed in mouse (Figure 2B).

**Figure 2.**
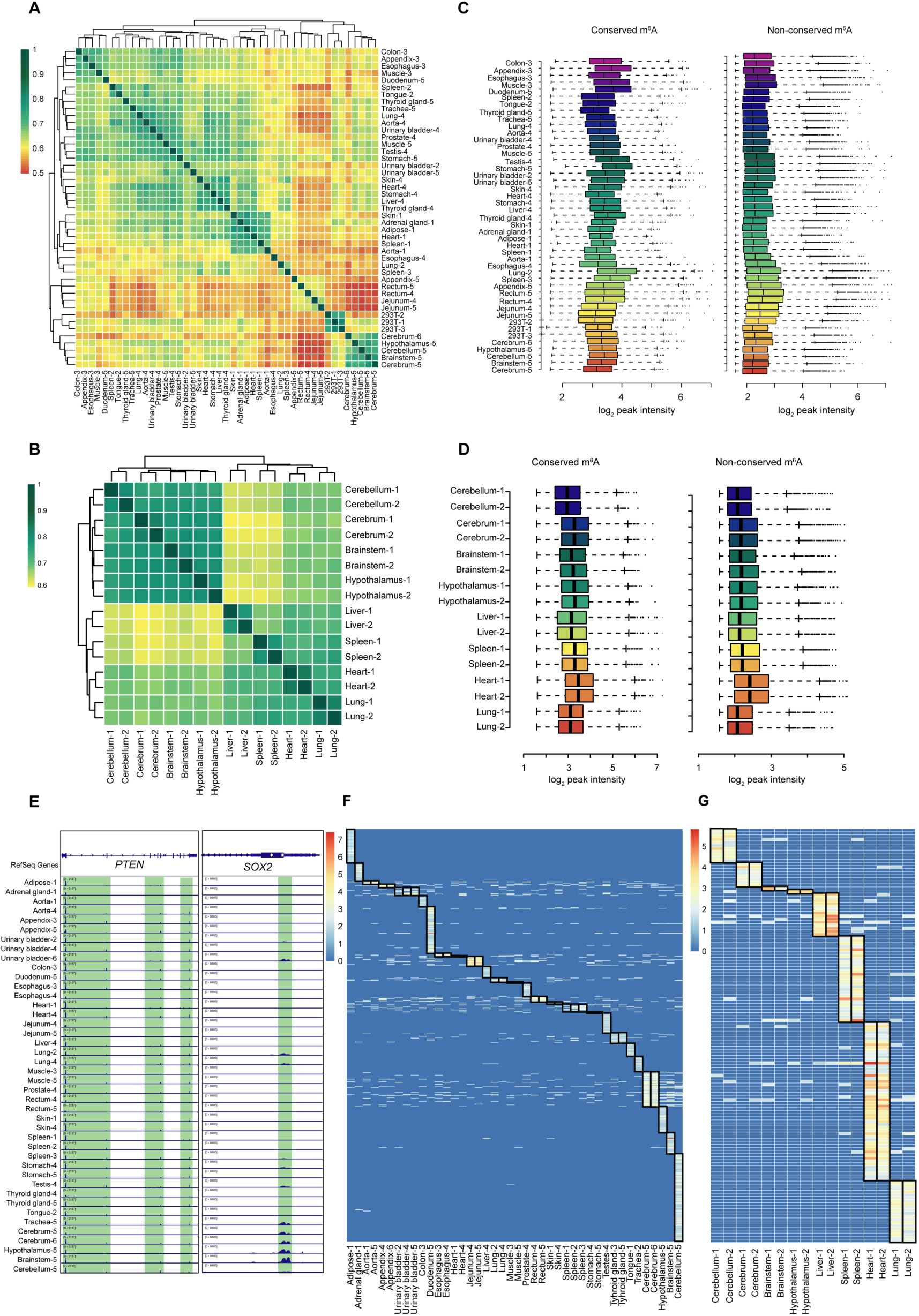
m^6^A methylome across different human and mouse tissues. (**A-B**) Heat map and dendrogram of Spearman correlations of the methylation levels across 42 human tissue samples and three HEK293T cell sapmles (**A**) and 16 mouse tissues (**B**) calculated using m^6^A intensity. The dendrogram was drawn based on the distance metric computed by the m^6^A intensity. The intensity of the color represents the similarity. Samples are denoted by the tissue name followed by a donor ID and an identical ID indicates that the tissue samples are from the same person or the same mouse. (**C-D**) Intensity of “conserved” and “non-conserved” m^6^A signals across various human tissues and HEK293T cell line (**C**) and mouse tissues (**D**). (**E**) Representative IGV views of conserved m^6^A peaks (*PTEN*) and non-conserved m^6^A peaks (*SOX2*). Green color denotes m^6^A signal. (**F-G**) m^6^A signals on ubiquitously expressed transcripts in all human tissues (**F**) or mouse tissues (**G**). The regions marked with boxes denote “tissue-specific” m^6^A signal.

We then assessed the intensity of the “conserved m^6^A peaks” (peaks identified in all tissues) and “non-conserved m^6^A peaks”. We found that the intensity of the conserved m^6^A peaks is significantly higher than that of the non-conserved m^6^A both in human and mouse (*P*<2.2*e^-16^) (Figures 2C and 2D), suggesting important regulatory roles of the conserved m^6^A signal. Take *PTEN* as an example for the conserved m^6^A peaks: it is stably expressed in all examined tissues and similarly m^6^A modified. Three m^6^A peak clusters which are located at 5’-UTR, CDS and around stop codon respectively can be identified in all tissues, indicating that m^6^A may play a generic regulatory role for *PTEN* (Figure 2E). Another gene, *SOX2*, was taken as an example for the non-conserved m^6^A peaks (Figure 2E).

Although we did not observe strong tissue-specificity of m^6^A methylome across different non-brain tissues in human and mouse, we nevertheless analyzed the tissue-specific m^6^A peaks among the tissues. Within the 9,431 ubiquitously expressed genes, we identified 21,480 m^6^A peaks, out of which 1,898 peaks are conserved across tissues while 594 m^6^A peaks are tissue-specific. Interestingly, all the tissue samples can be readily separated based on the small group of tissue-specific m^6^A signals for both human and mouse (Figures 2F and 2G). Moreover, we found that genes encoding transcripts with brain-specific m^6^A signals are enriched in head development functions (Figure S2E).

Besides mRNA, m^6^A is also found in lncRNA (Bohnsack and Sloan, 2018; Fu et al., 2014). Specifically, among the 42,873 lncRNAs expressed in the tissue samples, we totally identified 78,789 m^6^A peaks. In addition, we found 1,816 ubiquitously expressed lncRNAs but the number of conserved m^6^A peaks is limited (∼383). Similar to that of mRNA (Figure 2A), the methylome of ubiquitously expressed lncRNAs in brain tissues still show high tissue-specificity (Figure S2F). Moreover, we found that the intensity of the conserved m^6^A is significantly higher than that of the non-conserved m^6^A in human lncRNA (*P*<2.2*e^-16^) (Figure S2G).

### Tissue-specificity of m^6^Am in human and mouse

We next investigated m^6^Am profiles using hierarchical clustering based on the correlation among human tissues (the genes should be ubiquitously expressed in all tissue types). Similar to m^6^A, m^6^Am signals of the brain tissues were clearly resolved from that of the non-brain tissues (Figure 3A), suggesting that m^6^Am may be involved in the regulation of brain-related physiological processes. One example of conserved (*ZNHIT6*) and non-conserved (*AHCYL1*) m^6^Am peak is shown, respectively (Figure 3B). Moreover, we analyzed the m^6^Am methylome across 16 mouse tissues. Again, the different brain regions clustered together and are separated from non-brain tissues (Figure 3C). In fact, the m^6^Am signals of the mouse brain tissues appear to be further separated away from the rest of the tissues when comparing to the m^6^A clustering in Figure 2B.

**Figure 3.**
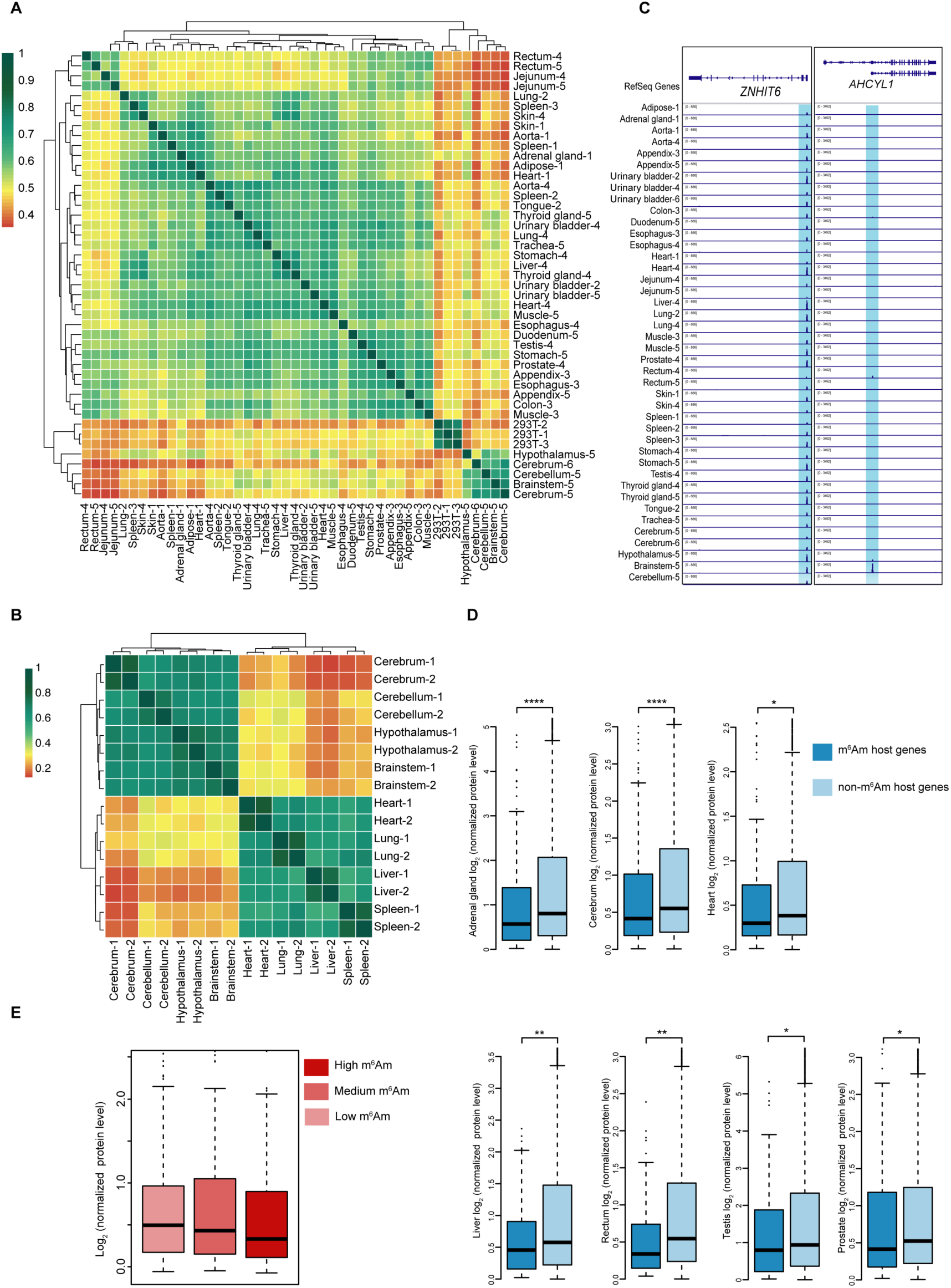
m^6^Am methylome across diverse human and mouse tissues. (**A-B**) Heat map and dendrogram of Spearman correlations of the m^6^Am levels across 42 human tissue samples and three HEK293T cells (**A**) and 16 mouse tissues (**B**). The dendrogram was drawn based on the distance metric computed by the m^6^Am intensity. The intensity of the color represents the degree of similarity. (**C**) Representative IGV views of conserved m^6^Am peaks (*ZNHIT*6) and non-conserved m^6^Am peaks (*AHCYL1*). Blue color denotes m^6^Am signal. (**D**) m^6^Am is negatively correlated with protein level in human tissues including adrenal gland, cerebrum, testis, heart, prostate, liver and rectum. Dark blue indicates the normalized protein level of m^6^Am-containing genes while the light blue represents the protein level of non-m^6^Am containing genes. The human proteome data was taken from a published study (see Method Details). Statistical significance of the difference was determined by Student’s t*-*test. ****p<0.0001; ***p < 0.001; **p < 0.01; *p < 0.05. (**E**) Boxplot showing that the normalized protein level gradually decreases with the increase of the m^6^Am signal.

### m^6^Am negatively correlates with protein level

We next tried to examine potential functions of m^6^Am in mRNA. We found that the m^6^Am mark is in general negatively correlated with protein level across the human tissues. For instance, the normalized protein level of m^6^Am marked genes is significantly lower than that of non-m^6^Am marked genes in multiple tissues including adrenal gland, cerebrum, heart, prostate, liver, rectum and testis (Figure 3D). A similar pattern can also be seen in lung, colon and esophagus, although the difference is not statistically significant owing to the limited number m^6^Am-marked genes under our strict cutoff for m^6^Am identification (Figure S3A). Moreover, we divided the m^6^Am marked genes into three sets on the basis of their modification level. Consistent with our previous finding, the normalized protein level gradually decreases with the enhancement of the m^6^Am signal (Figure 3E).

As the biological function of m^6^A modification relies on its reader proteins, it would be of great importance to identify potential m^6^Am readers that can directly regulate RNA metabolism. We adopted a computational pipeline to screen potential m^6^Am binding proteins using the m^6^Am modification signal identified in our study and the published crosslinking and immunoprecipitation followed by high-throughput sequencing (CLIP-seq) of 171 RNA binding proteins (RBPs) (Huang et al., 2018; Zhu et al., 2019). The top 15 and bottom 15 of the 171 RBPs are shown (Figure S3B). Interestingly, the top candidate, GNL3, is a known RBP that binds 5’UTR of mRNA; in addition, it does not bind m^6^A in various systems (Edupuganti et al., 2017). However, because the current sequencing resolution does not allow us to definitely distinguish m^6^Am from a nearby m^6^A in the 5’-UTR, future experiments are needed to test the specificity of the candidates in recognizing m^6^Am.

### Correlation between m^6^A & m^6^Am and the expression of writers & erasers

The extent to which variation in m^6^A and m^6^Am modification may be attributed to the expression of each methyltransferase component and demethylase remains unknown. By analyzing the peak numbers of m^6^A and m^6^Am in different tissues and the expression of the corresponding proteins, we found that the expression of methyltransferase components including *METTL3* and *WTAP* are positively correlated to the m^6^A variation and explains about 36.2% and 42.6%, of the m^6^A variation in human, respectively (Figures 4A and 4B). *METTL14* shows a weak correlation with m^6^A variation (Figure 4C). With regards to m^6^A erasers, the expression of *ALKBH5* is negatively correlated with m^6^A signal and explained approximately 38.2% of the variation in human (Figure 4D). In mouse, the expression of methyltransferase components including *METTL3* and *METTL14* are positively correlated to the m^6^A variation and explains about 50% and 68.1% of the variance while *WTAP* shows no correlation (Figures 4H-4J). *ALKBH5* is again negatively correlated with m^6^A signal and explained 88.6% of the m^6^A variance (Figure 4K). Therefore, despite slight difference, m^6^A methyltransferase components and demethylase generally contribute to the m^6^A abundance in human and mouse.

**Figure 4.**
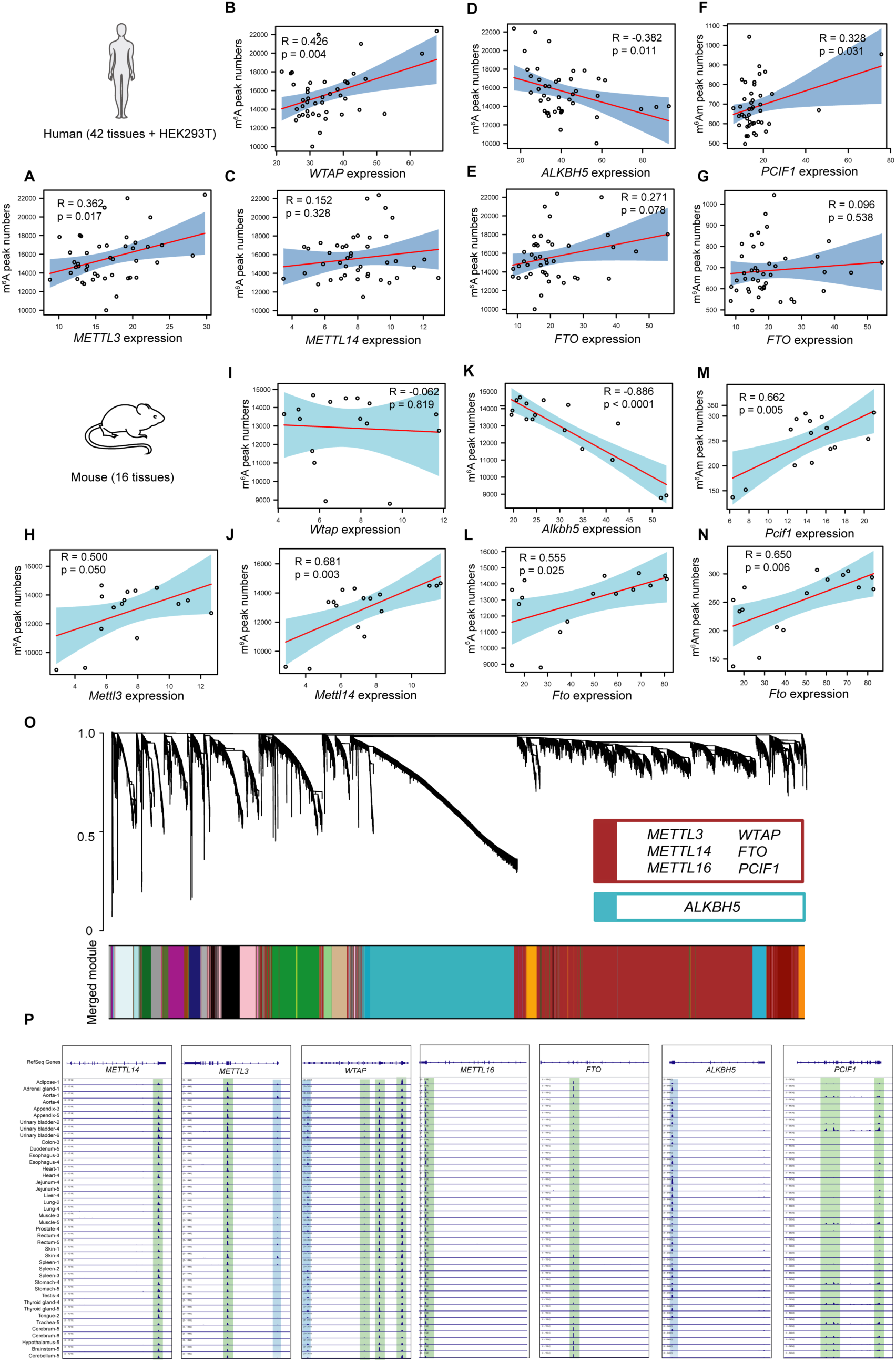
The correlation between m^6^A/m^6^Am and their corresponding writers & erasers. (**A-C**) Scatter plots showing the correlations between m^6^A peak numbers and the expression levels of m^6^A “writer” components including *METTL3* (**A**), *WTAP* (**B**) and *METTL14* (**C**) (quantified as the number of RNA-seq reads per kilobase of transcript per million mapped reads (RPKM)) across human tissues and HEK293T cell. (**D-E**) Correlations between m^6^A peak numbers and the expression levels of m^6^A erasers including *ALKBH5* (**D**) and *FTO* (**E**) across human tissues and HEK293T cell. (**F-G**) Correlations between m^6^Am peak numbers and the expression levels of *PCIF1*(**F**) and *FTO* (**G**) across human tissues and HEK293T cell. (**H-J**) Scatter plots showing the correlations between m^6^A peak numbers and the expression levels of *Mettl3* (**H**), *Wtap* (**I**) and *Mettl14* (**J**) across mouse tissues. (**K-L**) Correlations between m^6^A peak numbers and the expression levels of m^6^A erasers including *Alkbh5* (**K**) and *Fto* (**L**) across mouse tissues. (**M-N**) Scatter plots showing the correlations between m^6^Am peak numbers and the expression levels of *Pcif1*(**M**) and *Fto* (**N**) across mouse tissues. (**O**) Hierarchical cluster tree showing co-expression mRNA modules identified using WGCNA. Modules corresponding to mRNAs are labeled by colors. (**P**) IGV views showing the methylation marks on the transcripts of m^6^A & m^6^Am writers (METTL3, WTAP, METTL14, METTL16 and PCIF1) and erasers (FTO and ALKBH5). Green box indicates the m^6^A signal; blue box indicates the m^6^Am signal.

We also analyzed the extent to which variation of m^6^Am modification could be attributed to the expression of its writer and eraser. Very recently, the writer of m^6^Am has been identified by us and other labs independently: both in vivo and in vitro evidence has demonstrated that PCIF1 is specific for the cap-related, terminal m^6^Am modification (Akichika et al., 2018; Sun et al., 2018). The expression of PCIF1 accounts for approximately 32.8% and 66.2% of the variation in human and mouse, respectively (Figures 4F and 4M). FTO demethylates both m^6^A, m^6^Am and m^1^A, but its relative activity is recently shown to be dependent on its sub-cellular localization (Wei et al., 2018). While the localization of FTO in the human tissues was not examined, unexpectedly, we observed a non-negative correlation between FTO expression and m^6^A & m^6^Am variation, respectively (Figures 4E, 4G, 4L and 4N). To look into this non-negative correlation, we used GTEx datasets to perform co-expression analysis and found that FTO is highly co-expressed with both the m^6^A and m^6^Am methyltransferase components (Figure 4O). Considering that m^6^A and m^6^Am are reversible modifications involved in many different biological processes, such co-expression of methyltransferase components and demethylases could be beneficial to the dynamic regulation.

Notably, the transcripts of m^6^A and m^6^Am writers and erasers also contain these methylation marks (Figure 4P). For instance, we found conserved m^6^A peaks in *METTL3*, *METTL14*, *WTAP*, *METTL16*, *FTO* and *ALKBH5* across all human tissues and conserved m^6^Am peaks in *WTAP* and *ALKBH5*. We also identified alternative m^6^Am signals in *METTL3* and alternative m^6^A signals in *PCIF1*. Thus, the transcripts of the modification machineries for m^6^A and m^6^Am are also susceptible to epitranscriptomic regulation so that the crosstalk between m^6^A and m^6^Am is worth further explored.

### m^6^A and m^6^Am peaks are enriched for SNPs that are associated with diseases in human tissues

About millions of SNPs have been identified across multiple human genomes and the SNPs within m^6^A or m^6^Am peak regions are defined as m^6^A-related or m^6^Am-related SNPs. Here, we sought to investigate the relationship between m^6^A and SNPs across human tissues. Firstly, we sought to identify the relative distribution of m^6^A and SNPs. We found that m^6^A-containing regions are enriched for SNPs and the number of SNPs decreases as the distance from the m^6^A sites increases (Figure 5A). Within the m^6^A peaks (approximately 200-300nt), we observed ∼8,193-31,363 SNPs for each tissue, with jejunum containing the most abundant SNPs. In fact, non-brain tissues are more enriched for SNPs than brain tissues in the m^6^A peak regions (*P* < 2.2*e^-16^) (Figure 5B). To exclude the possibility that the observed enrichment is due to the position background, we also used the regions surrounding stop codon of the overall transcripts (position background control) or genome background for comparison. We found that the number of SNPs in m^6^A regions are still significantly higher than that of position background control and genome background (*P* < 2.2 *e^-16^) (Figure 5B).

**Figure 5.**
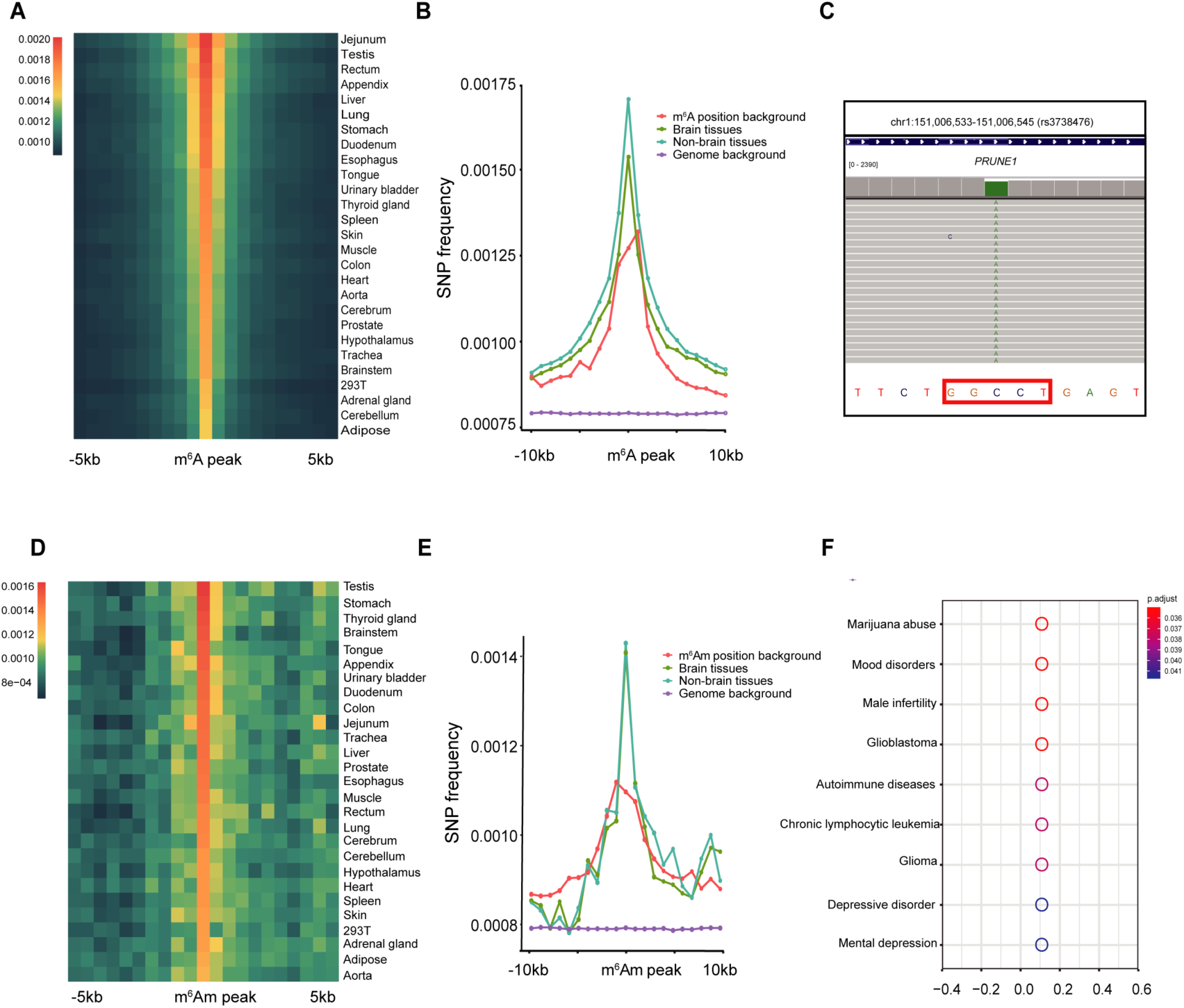
m^6^A and m^6^Am signals are enriched for SNPs. (**A**) Heat map showing the enrichment of SNPs flanking the m^6^A regions across human tissues. (**B**) SNP frequency in brain-related tissues (green line) and non-brain tissues (blue line) flanking the stop codon region (*P* <2.2*e-^16^). P-values were calculated using unpaired two-sided Mann-Whitney U-test. SNP frequency of the position background control (defined as the regions surrounding stop codon of the overall transcripts; red line), and genome background (defined as the regions which are selected randomly on the genome; purple line) are also shown. (**C**) A representative view of C-to-A mutation in the CDS region of *PRUNE1*, leading to the *de novo* generation of a GGACU consensus that is further m^6^A methylated in aorta. (**D**) Heat map showing that the m^6^Am regions are enriched for SNPs across human tissues. (**E**) SNP frequency in brain tissues (green line) and non-brain tissues (blue line) surrounding TSS regions (transcription start site) (*P* <2.2*e-^16^). P-values were calculated using unpaired two-sided Mann-Whitney U-test. SNP frequency of the position background control (defined as the regions surrounding TSS of the overall transcripts; red line) and genome background (defined as the regions which are selected randomly on the genome; purple line) are also shown. (**F**) Top Disease ontology (DO) categories of the m^6^Am-related SNPs in human brain tissues including cerebellum, cerebrum, hypothalamus and brainstem.

We next analyzed the m^6^A-related SNPs and found that the number of synonymous and nonsynonymous SNPs are comparable, which accounts for 48.9% and 49.7% across all tissues, respectively (Figure S4A). In addition, m^6^A-related SNPs are mainly located at exon (>90%), among which 48.4% and 49.2% fall within the 3’-UTR and CDS, respectively (Figure S4B and S4C). Interestingly, m^6^A-related SNPs also include mutations that lead to gain of m^6^A modification (or “m^6^A-gain variants”). Take *PRUNE1* as an example: a synonymous mutation, C-to-A mutation (rs3738476) in the CDS region generates a GGACU sequence *de novo*, which enables this site to be m^6^A modified (Figure 5C). In total, we identified ∼0.76% m^6^A-related SNPs to be within a 16nt window of a putative RRACH consensus motif. We further used “Disease ontology (DO)” to reveal the relevance of m^6^A-related SNPs in diseases, and found that the m^6^A-related SNPs are highly correlated with diseases including colorectal cancer, colorectal carcinoma and coronary artery diseases (Figure S4D).

We next assessed the correlation between m^6^Am and SNPs. After ruling out potential positional and genomic background (*P* < 2.2 *e-^16^), we found that the m^6^Am regions are also enriched for SNPs in human tissues (Figure 5D). Different from m^6^A, the SNP frequency has no significant difference between brain tissues and non-brain tissues (*P*=0.063) (Figure 5E). We next analyzed m^6^Am-related SNPs and found that the number of synonymous and nonsynonymous SNPs accounts for 45.5% and 53.1% across all tissues, respectively (Figure S4E). In addition, m^6^Am-related SNPs are mainly located at exon (>90%), among which 51.6% and 42.5% fall within the 5’-UTR and CDS, respectively (Figures S4F and S4G). We further revealed that m^6^Am-related SNPs appear to be associated with asthma (Figure S4H). Interestingly, when we analyzed m^6^Am related SNPs in brain tissues, we found that they are specifically enriched for brain-related diseases including marijuana abuse, mood abuse and glioblastoma (Figure 5F). Together, our results suggest a connection between epitranscriptomic marks and risk for diseases.

### m^6^A peaks are enriched at the microRNA target sites

Because m^6^A peaks are enriched near the stop codon and in the 3’-UTR and microRNA (miRNA) target sites are frequently observed within 3’-UTR, we next sought to investigate whether m^6^A are associated with miRNA binding sites. We found that more than 77%-80% m^6^A-containing transcripts have at least one miRNA binding site. Further analysis revealed that nearly all the m^6^A peaks (>96%) could pair with miRNAs with relatively strict alignment criteria across all tissues. Regions surrounding the m^6^A peaks contain ∼9,786-14,104 miRNA binding sites, with cerebellum m^6^A regions contain the most miRNA target sites. In all the tissues, microRNA targeting sites show an enriched distribution around the m^6^A peaks (Figure S5A). To rule out that the enrichment of the miRNA targets is due to potential positional effects, we analyzed the miRNA targets near stop codon and found that the distribution pattern of miRNA near m^6^A is significantly different from the pattern near stop codon (Figure S5B).

Among the m^6^A consensus motif “RRACH”, “GGACH” is the most frequent in all tissues (Figure S5C). We used relatively strict criteria in which at most 1nt mismatch was allowed, and found that m^6^A peaks could be targeted by ∼ 423 miRNAs. For instance, the strongest, GGACH consensus motif was inversely complementary to the seed region of many miRNAs. To further explore the relationship between microRNA and m^6^A-containing genes, we next searched miRNAs that are indeed expressed in the corresponding tissues. We found that 78 miRNAs were stably expressed in all tissues, of which, 11 could pair with m^6^A motif (Figure S5D). We also identified tissue-specific miRNAs across urinary bladder, brain, liver, lung and testis, and revealed that they are inversely complementary to the m^6^A motifs as well (Figure S5E). Finally, the miRNAs specifically expressed in brain tissues are most likely to target m^6^A peaks: about 27% of brain-specific miRNAs could pair with m^6^A.

### m^6^A and m^6^Am methylome conservation across species

To demonstrate the conservation of m^6^A between human and mouse, we first compared the m^6^A methylome in human cerebellum with that of mouse cerebellum. We observed that the m^6^A-containing orthologous genes in the cerebellum of the two species exhibit a high overlap (Figure 6A), consistent with the comparative analysis of m^6^A methylome using HepG2 cell lines and mouse liver (Dominissini et al., 2012). However, the degree of conservation of m^6^A between tissues and species was still unknown. Intriguingly, although the overlap between matched tissues from different species is already high (Figure 6A), all tissues of same species demonstrate higher similarity of overall m^6^A signals and tend to group together than the same tissues of different species (Figure 6B and S6A). Moreover, we divided the orthologous genes into 5’-UTR, CDS and 3’-UTR segments and again observed the similar clustering of m^6^A methylome by species (Figures S6B-6D). In addition, we also picked out the housekeeping genes as reported before (Eisenberg and Levanon, 2013) and found that the overlap of the m^6^A-containing housekeeping genes in different tissue of the same species, for instance human cerebellum and heart, is significantly higher than that of the matched tissue from different species, for instance human and mouse cerebellum (*P* < 2.2 *e^-16^) (Figures 6C and 6D), implying that the m^6^A methylome of housekeeping genes contributed to the overall clustering pattern.

**Figure 6.**
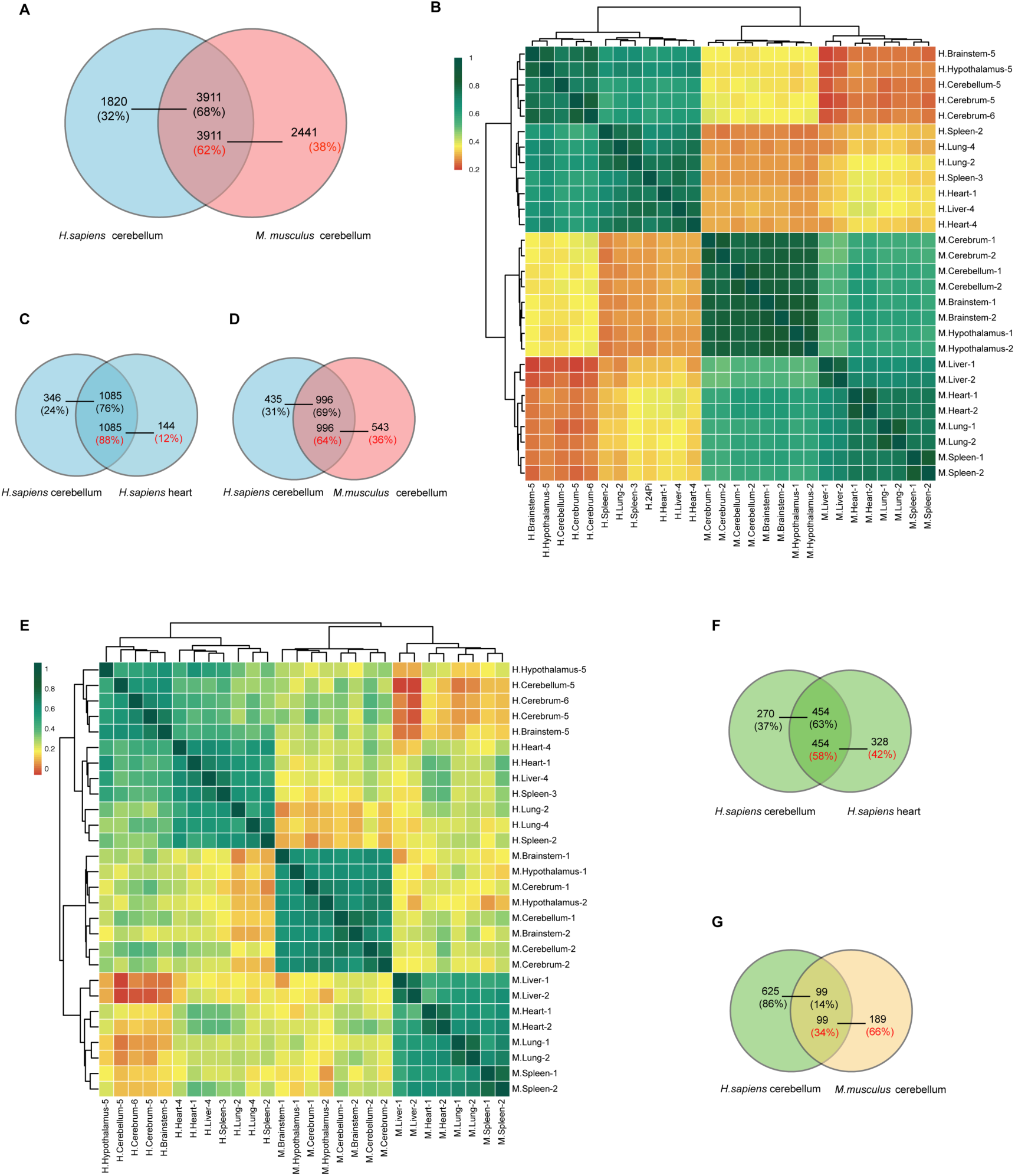
Comparison of m^6^A methylome between human and mouse. (**A**) Venn diagram showing the overlap of m^6^A-containing orthologous genes across human cerebellum and mouse cerebellum. (**B**) Heat map and dendrogram of Spearman correlations of the m^6^A levels of the matched tissues between human and mouse. (**C-D**) Venn diagram showing the overlap of m^6^A-containing housekeeping genes between human cerebellum and human heart (**C**), and between human cerebellum and mouse cerebellum (**D**). (**E**) Heat map and dendrogram of Spearman correlations of the m^6^Am levels of the matched tissues between human and mouse. (**F-G**) Venn diagram showing the overlap of m^6^Am-containing orthologous genes between human cerebellum and human heart **(F**), and between human cerebellum and mouse cerebellum (**G**).

To obtain a comparative view of m^6^Am methylome across human and mouse tissues, we performed m^6^Am analysis and again, observed that different tissue samples were largely grouped by species rather than tissue types (Figures 6E and S6E). Take the human cerebellum, human heart and mouse cerebellum as examples, the m^6^Am-containing orthologous genes in the human cerebellum and heart shows significantly higher overlap than that between the human cerebellum and mouse cerebellum (*P*<2.2*e^-16^) (Figures 6E and 6G).

## DISCUSSION

Despite their importance in RNA biology, m^6^A and m^6^Am modifications were not interrogated across diverse human and mouse tissues. In this study, we measured the level of m^6^A and m^6^Am and provide the first transcriptome-wide characterization of m^6^A and m^6^Am methylome in human and mouse tissues. We show that the overall m^6^A and m^6^Am methylome are similar across various non-brain tissues, whereas brain tissues are highly specific. Interestingly, a small subset of tissue-specific m^6^A signals nevertheless distinguish different human and mouse tissue types. We also find that m^6^A and m^6^Am are partially associated with their writers and erasers. We uncover associations between m^6^A & m^6^Am and SNPs, and found a negative correlation between m^6^Am and protein expression. Finally, we show that species is a stronger factor than tissue type in determining the m^6^A and m^6^Am methylome. Collectively, our study reveals that m^6^A and m^6^Am are widespread and dynamically regulated epitranscriptomic marks across human and mouse tissues.

Our transcriptome-wide mapping relies on an antibody that binds to both m^6^A and m^6^Am. The signals of m^6^A and m^6^Am are not directly separated by the sequencing method, but are rather differentiated bioinformatically by using a variety of criteria as previously reported (Schwartz et al., 2014). Even though the two modifications reside in different biological context (internal vs terminal; and different consensus motifs), the bioinformatics pipeline can not definitely distinguish them. Nevertheless, our quantitative MS analysis relies on the different molecular weight of m^6^A and m^6^Am, and hence the modification levels across different human tissues are accurate. Notably, the number of m^6^A and m^6^Am peaks and the overall methylome we identified agrees well with the quantitative modification level by MS. While potential knock-out experiments of modification enzymes could be performed in cell lines to cleanly separate the two marks, it is currently not possible to remove methyltransferases in human tissues. Thus, it will be of great interest to have epitranscriptomic tools in future to directly and definitely distinguish m^6^A from m^6^Am.

Our analyses reveal that m^6^A and m^6^Am are highly correlative across non-brain tissues at the overall level, while that of brain tissues show high tissue-specificity. Such finding is reminiscent of previous studies of inosine and 5mdC (Schultz et al., 2015; Tan et al., 2017; Ziller et al., 2013). The inosine profiles across different tissues are also highly correlated, with the exception that brain regions can be resolved from non-brain tissues by principal component analysis; while for 5mdC, nearly 80% CpG sites are similarly methylated and the differentially methylated regions (DMRs) only accounts for 20% CpG sites in diverse cell lines (Schultz et al., 2015; Ziller et al., 2013). Nevertheless, we still uncovered many transcripts that exhibit tissue-specific m^6^A methylation patterns, demonstrating that m^6^A signal can be used to classify different types of human and mouse tissues, consistent with the use of DMRs to classify unknown samples and identify representative signature regions that recapitulate major DNA methylation dynamics. As for the observed tissue-specificity of m^6^A and m^6^Am in brain tissues, many studies have reported the importance of m^6^A modification in human brain development and neurological disorders (Du et al., 2018). We also highlight the specificity of both m^6^A and m^6^Am in brain tissues and show the brain-specific m^6^A are enriched in brain-related function, further supporting that m^6^A could play unique roles for brain tissues. It’s worth mentioning that during the preparation of this manuscript, m^6^A methylome in human fetus tissues was reported (Xiao et al., 2019). Although different human tissue samples were used, it appears that m^6^A methylome in brain tissues of fetus is also highly specific.

m^6^Am is recently shown to be a reversible epitranscriptomic mark (Mauer et al., 2017; Wei et al., 2018). However, the ability of FTO to demethylate both m^6^A and m^6^Am renders it difficult to functionally separate the two modifications. More recent identification and characterization of the m^6^Am writer protein PCIF1 by us and others have shown that PCIF1 is specific for the cap-related m^6^Am but not internal m^6^A, providing an orthogonal system to specifically dissect the biological roles of m^6^Am (Akichika et al., 2018; Sun et al., 2018). In this study, we show that the expression of PCIF1 is correlated with m^6^Am. In addition, we found that m^6^Am is negatively correlated with protein level, providing insights into the functional roles of m^6^Am. It would also be interesting to integrate such m^6^Am methylome data to identify potential m^6^Am reader proteins in the future.

Our profiling reveals greater m^6^A and m^6^Am difference between species than tissue types, which is highly suggestive of stronger cis-directed regulation of m^6^A and m^6^Am methylome. This is also observed for inosine, for which tissue samples are also largely grouped by species rather than by tissue type. Additionally, it is found that inosine sites edited similarly between species have more conserved flanking sequences than sites edited differentially. Once more m^6^A and m^6^Am datasets (ideally high resolution methylome) in other species become available in future, the influence of flanking sequences on m^6^A and m^6^Am signal can also be investigated. Besides, the regulation of m^6^A and m^6^Am by species-specific trans-acting factors is also worth exploration. For instance, a novel trans-regulatory factor AIMP2, which enhances the degradation of the ADAR proteins, has been identified for inosine (Tan et al., 2017). Future studies aiming at uncovering additional cis- and trans-regulators of m^6^A and m^6^Am are necessary to determine how precise control of methylation is achieved in a myriad of biological contexts.

In summary, our work reveals the dynamic landscape of m^6^A and m^6^Am across human tissues and provides a resource for functional studies of m^6^A and m^6^Am in the future.

## ACKNOWLEDGMENTS

The authors would like to thank Hu Zeng, Bo He and Xiaoxue Zhang for experiments; Xiaoyu Li and Dongsheng Bai for discussions. We acknowledge Shanghai Epican Genentech Co. Ltd. for helping the sample collection process. This work was supported by the National Natural Science Foundation of China (nos. 2182570 and 91740112 to C.Y.; 81602513 to J. C.), the Joint Laboratory of International Scientific and Technological Cooperation (Beijing Municipal Science and Technology Commission, Cooperative study of RNA modification detection technology and early diagnosis of disease).

## AUTHOR CONTRIBUTIONS

J.L. and C.Y. conceived the project, designed the experiments and wrote the manuscript. J.L. performed all experiments. K.L., J.L. and X.Z. designed and performed the bioinformatics analysis with the help of X.X., H.M. and C.Y. J.C., M.Z., Z.H., X.X., and J.F collected human and mouse tissue samples. All authors commented on and approved the paper.

## STAR METHODS

### CONTACT FOR REAGENT AND RESOURCE SHARING

Further information and requests may be directed to, and will be fulfilled by the corresponding author Chengqi Yi.

### EXPERIMENTAL MODEL AND SUBJECT DETAILS

HEK293T was maintained at 37 °C in DMEM medium (Gibco) supplemented with 10% FBS (Gibco) and 1% penicillin/streptomycin (Gibco).

### METHOD DETAILS

#### Human tissues

42 different somatic tissue samples were collected from six post-mortem healthy Chinese including 5 female donors and 1 male donor. All samples were obtained with informed consent under a protocol approved by the ethics committee of School of Basic Medical Sciences of Fudan University, China.

#### Mouse tissues

16 mouse tissue samples were collected from 2 male C57BL/6J mice and used at 7 weeks of age. All mice were bred and kept under specific pathogen-free conditions in the Laboratory Animal Center of Peking University in accordance with the National Institute of Health Guide for Care and Use of Laboratory Animals. Animal protocols were approved by the Institutional Animal Care and Use Committee at Peking University.

#### Cell culture

HEK293T cells was cultured in DMEM medium (Gibco) supplemented with 10% (v/v) FBS (Gibco) and 1% penicillin/streptomycin (Gibco, 15140) at 37 °C with 5% CO_2_.

#### RNA extraction and DNase treatment

Total RNA was extracted from tissues using TRIzol reagent (Invitrogen,15596018), followed by DNase I (NEB, M0303L) treatment to remove DNA contamination. Additional phenol-chloroform isolation and ethanol precipitation treatment was performed to remove enzyme contamination.

#### Quantitative analysis of m^6^A and m^6^Am level in different tissues

For the quantification of m^6^A and m^6^Am level in tissues, 150 ng purified RNA was decapped with 10 U RppH (NEB, M0356S) in ThermoPol buffer for 5 hours at 37 ℃. RNA was further digested by 1U nuclease P1 (Sigma, N8630) in 20 μl buffer containing 10 mM NH4Ac, pH 5.3 at 42 ℃ for 3 h. Subsequently, 1U rSAP (NEB, M0371S) and 5 μl 0.5 M MES buffer, pH 6.5 were added and the mixture was incubated at 37 ℃ for additional 6 h. The digested RNA was injected into a LC-MS/MS which includes the ultra-performance liquid chromatography with a C18 column and the triple-quadrupole mass spectrometer (AB SCIEX QTRAP 5500). The m^6^A and m^6^Am level were detected in the positive ion multiple reaction-monitoring (MRM) mode and was quantified by the nucleoside to base ion mass transitions (282.0-to-150.1 for m^6^A, 296.0-to-150.1 for m^6^Am, 268.0 -to- 136.0 for A). The concentrations of m^6^A and m^6^Am level in human tissues were calculated from the standard curve which was generated from pure nucleoside standards.

#### m^6^A-seq with low input RNA across human and mouse tissues

This procedure was performed according to the recently described “Refined RIP-seq” with several modifications(Zeng et al., 2018). 3 μg total RNA was fragmented into ∼130-nucleotide-long fragments by magnesium RNA fragmentation buffer (NEB, E6150S). The fragmentation was stopped by adding RNA fragmentation stop solution followed by ethanol precipitation. 8 ng of fragmented total RNA was used as input and remained RNA was used to do the m^6^A-seq. Briefly, RNA was denatured at 65 °C for 5 min, followed by chilling on ice. 30 μl protein A magnetic beads (Thermo, 10002D) and 30 μl protein G magnetic beads (Thermo, 10004D) were mixed together and washed twice using the IPP buffer (10 mM Tris-HCl, pH 7.5,150 mM NaCl, 0.1% IGEPAL CA-630) and resuspended in 500 μl of IPP buffer. The 6 μg of affinity purified anti-m^6^A polyclonal antibody (Millipore, ABE572) was added to the beads and incubated at 4 ℃ for about 6 h. After the beads-antibody incubation, the beads were washed twice by IPP buffer and resuspended with 500 μl mixture which contains 100 μl of 5×IPP buffer, fragmented total RNA, and 5 μl of RNasin Plus RNase Inhibitor (Promega, N2615) and incubated at 4 ℃ for another 2 h. The beads-antibody-RNA mixture was washed with twice IPP buffer, twice with low-salt IP buffer (10 mM Tris-HCl, pH 7.5, 50 mM NaCl, 0.1% IGEPAL CA-630), and twice high-salt IP buffer (10 mM Tris-HCl, pH 7.5, 500 mM NaCl,0.1% IGEPAL CA-630). After extensive washing, bound RNA was eluted from the beads with 6.7mM N6-methyladenosine (Sigma-Aldrich, M2780) in IPP buffer and additional phenol-chloroform isolation and ethanol precipitation treatment was performed to purify the RNA. Fragmented total RNA (Input) and immunoprecipitated RNA (IP) were subjected to library construction using SMARTer® Stranded Total RNA-Seq Kit v2 - Pico Input Mammalian (634413, Takara – Clontech, Japan) according to the manufacturer’s protocol. Reverse transcription was performed using random primers and the ribosome cDNA (cDNA fragments originating from rRNA molecules) after cDNA synthesis using probes specific to mammalian rRNA. Libraries for immunoprecipitated RNA were PCR amplified for 14 cycles whereas 11 cycles were used for input RNA. The libraries were sequenced on Illumina Hiseq X10 with paired-end 2X 150 bp read length.

#### Reads pre-processing and alignment

Raw sequencing reads were firstly subjected to Trim_galore (http://www.bioinformatics.babraham.ac.uk/projects/trim_galore/) for quality control and trimming adaptor. The quality threshold was set to 20, and the minimum length required for reads after trimming was 30 nt. All reads that mapped to human rRNA by TopHat2 (version 2.0.13)(Kim et al., 2013) were removed. Processed reads were mapped to genome (hg19, UCSC Genome Browser and mm10, UCSC Genome Browser) using HISAT2 (version 2.1.0)(Kim et al., 2015) with default parameters, and separated by strand with in-house scripts.

#### Identification of putative m^6^Am and m^6^A sites

For genome-base peak caller MACS2 (version 2.1.1)(Feng et al., 2012), the effective genome size was set to 2.7*10^9^ for human and 1.87*10^9^ for mouse, under the option of *-nomodel* and *p-value* cutoff 0.01, and all input bam files were retained the same reads number. Peak annotated by annotatePeaks.pl (Homer version 4.8)(Heinz et al., 2010), and peaks reads coverage were showed by IGV (version 2.4.15)(Thorvaldsdottir et al., 2013). An m^6^Am peak was identified when a peak region contains an adenosine at the transcription start site, following a previously published procedure (Schwartz et al., 2014). The adenosine transcription start site was defined as: the number of reads starting at 100 nt before or after the annotated transcription start site (TSS) was greater than 5 in both IP and input samples, and an adenosine was at the detected site or at the position immediately preceding it. The m^6^A and m^6^Am peaks from all tissue samples were all merged to generate the reference peak list.

#### Peak intensity

The cover of a peak region was defined as its Reads Per Kilobase Million (RPKM) value, and then the peak intensity for the corresponding region was calculated as (IP cover) / (Input cover).

#### Tissue-specific m^6^A peaks

Peaks specifically identified in a particular tissue or the intensity of peaks in a particular tissue at least five times those in all other tissues.

#### Analysis of RNA-seq data

Paired-end, adapter-clean reads were mapped to human and mouse genome (hg19 and mm10, UCSC Genome Browser) using TopHat2 (version 2.0.13) with default parameters. The expression of transcripts was quantified as FPKM by Cufflinks (version 2.2.1)(Trapnell et al., 2010). Tissue conserved genes and tissue specific genes were identified using Summarized Experiment algorithm in TissueEnrich R package(Jain and Tuteja, 2018). Genes were classified into five categories based on their expression across 42 human tissues as previously published (Uhlen et al., 2015): (I) ubiquitously expressed genes: a large fraction (59%) of the genes that were detected in all analyzed tissues (FPKM>1); (II) tissue enhanced genes: genes with only a moderately elevated expression and mRNA levels in a particular tissue at least five times average levels in all tissues; (III) group-enriched genes: mRNA levels at least five times those in a small number of tissues (2-7); (IV) tissue enriched genes: mRNA levels in one tissue type at least five times the maximum levels of all other analyzed tissues; (V) mixed genes: detected in fewer than 42 tissues in human tissues in mouse but not elevated in any tissues.

#### Motif discovery and GO enrichment analysis

For the analysis of sequence consensus, the top 1000 peaks were chosen for de novo motif analysis with MEME (version 4.12.0)(Bailey et al., 2009), with taken 100 nt long peak summit centred sense sequences as input. Gene Ontology (GO) enrichment analyses were performed using DAVIED web-based tool (http://david.abcc.ncifcrf.gov/)(Huang et al., 2009).

#### GTEx data download and WGCNA analysis

Genotype-Tissue Expression (GTEx) project data (version 7th) were download from https://gtexportal.org/home/, which contain 500 RNA-seq sample. We perform weighted gene co-expression network analysis (WGCNA) with GTEx RNA-seq data. And the count of gene’s TPM (Transcripts Per Million) higher than 0 have to at least larger than 50. Meanwhile miRNA, snoRNA, snRNA and tRNA were removed from the gene list. For WGCNA in R (version 3.5)(Langfelder and Horvath, 2008), the soft power was set as 5 and merge the dynamic module with module distance cutoff was set as 0.3.

#### Protein expression analysis

Protein expression data was downloaded from Human Proteome Map (HPM) database (Kim et al., 2014) and protein expression level was normalized by gene expression level.

#### Relationship analysis of miRNA with m^6^A sites

Human miRNA Expression Database (miRmine) data were download from http://guanlab.ccmb.med.umich.edu/mirmine/, Tissue conserved miRNA and tissue specific miRNA were identified using Summarized Experiment algorithm. Human mature miRNA sequences were downloaded from miRbase (Release 22.1)(Kozomara and Griffiths-Jones, 2014). The miRNA seed region (5’ 2-8 nt) was obtained and compared with m^6^A peak region using Bowtie (version 1.1.2)(Langmead et al., 2009).

#### Non-coding RNA annotation

The human lncRNA annotation was download from LNCipedia (version 5.2)(Volders et al., 2018), https://lncipedia.org/download, while the snoRNA and snRNA annotation were download from NCBI RefSeq. Finally, all RNA annotations was merged to build RNA annotation file.

#### Comparison of m^6^A and m^6^Am sites between human and mouse

To compare human and mouse m^6^A site, Human and mouse orthologous gene set was downloaded from MGI (http://www.informatics.jax.org/), and only orthologous genes were used to analyze correlations.

#### Correlation analysis of SNPs with m^6^A and m^6^Am signal

To explore the correlation between SNP and m^6^A site, we extended 5 kb upstream and downstream of each m^6^A peak. Then the total 10 kb region was divided into 20 windows, with each window spanning 500 bp. The SNP database, which was obtained from 1000 Genomes Project (ftp://gsapubftp-anonymous@ftp.broadinstitute.org/bundle/hg19/1000G_omni 2.5.hg19.sites.vcf.gz), intersected with the m^6^A site peaks and the extended windows (bedtools v2.27.1) to count the SNP frequency (Quinlan and Hall, 2010). We next calculated the SNP frequency of transcripts that contain m^6^A-related SNPs and of the position background which are defined as regions flanking stop codon (∼400bp) to eliminate the influence of position background. Besides, we also selected 1,000,000 genome random regions from hg19 (random seed 6135). The SNP frequency of genome random regions are defined as genome background. The location of m^6^A-related SNP sites were annotated by the transcript segments, including the CDS, 3′ UTR, 5′ UTR, start codon, stop codon and etc., and the type of SNPs were annotated into synonymous and nonsynonymous by ANNOVAR (Wang et al., 2010). We used samtools(v1.7)(Li, 2011) to identify SNPs and m^6^A-related SNPs (Jiang et al., 2018) to get the m^6^A-gain variants from the m^6^A IP data. Lastly, we used DOSE(v3.2.0)(Yu et al., 2015) to analyze the correlation between m^6^A-related SNPs and diseases. Correlation analysis of SNP and m^6^Am was similar to that of m^6^A. It is noteworthy that we also defined and calculated the SNP frequency of position background (regions surrounding TSS) and genome background to ensure the m^6^Am enrichment is not false-positive.

#### Prediction of m^6^Am-binding proteins

We adopted a computational pipeline (Huang et al., 2018) to screen potential m^6^Am binding proteins using the m^6^Am modification signal identified in our study and the published crosslinking and immunoprecipitation followed by high-throughput sequencing (CLIP-seq) of 171 RNA binding proteins (RBPs) from POSTAR, which provides annotations and functions of RNA binding proteins (RBPs) as well as RBP binding sites (Zhu et al., 2019). The CLIP–seq peaks of each RBP were intersected with m^6^Am peaks identified in our study to calculate the ratio between the number of peaks overlapping with m^6^Am and the total number of RBP binding sites (m^6^Am-containing peak numbers/total RBP binding sites numbers). Besides, we also use the total m^6^Am peak numbers of the tissues to avoid false-positive. Moreover, the regions flanking the transcription start sites (from the transcript start site extend backward 400bp, referred to non-m^6^Am region) of non-m^6^Am harboring transcripts were selected and were intersected with the RBPs binding sites as negative control. We defined and calculated the RBP enrichment score (ES) as follows:

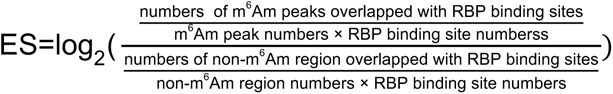

### QUANTIFICATION AND STATISTICAL ANALYSIS

p-values were calculated using unpaired two-sided student’s t test. ***p < 0.001; **p < 0.01; *p < 0.05; n.s., non-significant. As to SNPs analysis, p-values were calculated using unpaired two-sided Mann-Whitney U-test. ∗p < 0.05, ∗∗p < 0.01. N.S. stands for not significant. Error bars represent mean ± SD.

### DATA AND SOFTWARE AVAILABILITY

The raw sequence data reported in this paper have been deposited in the Genome Sequence Archive in BIG Data Center, Beijing Institute of Genomics (BIG), Chinese Academy of Sciences. A summary of the m^6^A and m^6^Am signal identified in human and mouse tissues and cell lines can be found in Table S1. All other data supporting the findings of this study are available from the corresponding author on reasonable request.

